# Identification of an A20 critical region harboring missense variations that lead to autoinflammation

**DOI:** 10.1101/2022.07.29.502017

**Authors:** Elma El Khouri, Camille Louvrier, Eman Assrawi, Alexandre Nguyen, William Piterboth, Samuel Deshayes, Alexandra Desdoits, Bruno Copin, Florence Dastot Le Moal, Sonia-Athina Karabina, Serge Amselem, Achille Aouba, Irina Giurgea

## Abstract

A20 haploinsufficiency (HA20) is an autoinflammatory disease caused by heterozygous loss-of-function variations in *TNFAIP3,* the gene encoding the A20 protein. Diagnosis of HA20 is challenging due to its heterogeneous clinical presentation and the lack of pathognomonic symptoms. While the pathogenic effect of *TNFAIP3* truncating variations is clearly established, that of missense variations is difficult to determine. Herein, we identified a novel *TNFAIP3* variation, p.(Leu236Pro), located in the A20 Ovarian Tumor (OTU) domain and demonstrated its pathogenicity. In the patients’ primary cells, we observed reduced A20 levels explained by enhanced degradation. We showed a disrupted ability of A20_Leu236Pro to inhibit the NF-κB pathway. Review of previously reported *TNFAIP3* missense variations revealed that only 3/7 are pathogenic. Through structural modeling we showed that the residues involved in OTU pathogenic missense variations establish common interactions. Interpretation of newly identified missense variations is challenging, requiring, as illustrated here, functional demonstration of their pathogenicity. Together with functional studies, *in-silico* structure analysis is a valuable approach that allowed us to unveil a region within the OTU domain critical for A20 function.

## INTRODUCTION

The zinc finger protein A20 is a ubiquitin-editing enzyme encoded by the *TNFAIP3 (tumor necrosis factor a-induced protein 3*) gene that contains an amino-terminal ovarian tumor (OTU) domain followed by seven zinc finger (ZnF) domains. Due to its E3 ligase activity (ZnF4), its deubiquitinase function (OTU) and its ubiquitin-binding domain (ZnF7), A20 regulates the fate of several substrates such as RIPK1 (Receptor-Interacting Serine/Threonine-Protein Kinase 1), NEMO (NF-Kappa-B Essential Modulator) and TRAF6 (TNF Receptor Associated Factor 6), all converging to the canonical NF-κB pathway (Boone et al., 2004; De et al., 2014; Tokunaga et al., 2012; Wertz et al., 2004). Owing to these various functions, A20 plays a pivotal role in the regulation of inflammation and immunity by inhibiting the canonical NF-κB signaling pathway and by preventing cell death (Lork et al., 2017; Martens and van Loo, 2020). Abnormal expression or function of A20 therefore contributes to the onset of several autoimmune or inflammatory disorders such as Crohn’s disease, systemic lupus erythematosus, rheumatoid arthritis, type-1 diabetes mellitus, psoriasis, and atherosclerosis as shown by genome-wide association studies (reviewed in (Malynn and Ma, 2019)). Furthermore, germline variations in *TNFAIP3* lead to an autoinflammatory disease of autosomal dominant inheritance referred to as A20 haploinsufficiency (HA20; MIM#616744) where insufficient suppression of NF-κB activity is observed (Zhou et al., 2016). HA20 is a Behçet-like autoinflammatory syndrome of earlier onset associated with variable clinical signs; the hallmark features of HA20 are recurrent fever, painful oral, genital and/or gastrointestinal ulcers with diarrhea, arthralgia or polyarthritis (Aeschlimann et al., 2018). Since the first description of HA20 (Zhou et al., 2016), more than 25 *TNFAIP3* truncating variations have been reported (reviewed in (Yu et al., 2020)) whereas the pathogenic significance has been confirmed for only a part of the 7 reported missense variations (Yu Chen et al., 2020; Dong et al., 2019; Kadowaki et al., 2021; Mulhern et al., 2019; Shigemura et al., 2016). Indeed, a body of genetic and experimental evidence is necessary to determine non-ambiguously the deleterious character of missense variations.

In this study, we report a non-previously described *TNFAIP3* missense variation, c.707T>C, p.(Leu236Pro), identified in three patients from the same family. We provide a phenotypic description of all affected patients, as well as cytokine and A20 level quantification in patient-derived samples. Furthermore, we assessed the pathogenicity of this variant through *in vitro* molecular and cellular assays. Interestingly, by studying the location and the amino acid interactions of Leu236 and of the other amino acids involved in pathogenic *TNFAIP3* missense variations, we identified a critical region within the OTU domain of A20 where the pathogenic *TNFAIP3* missense variations are in close proximity and the affected residues share common amino acid interactions on the 3D protein structure.

## RESULTS

### Clinical features of HA20 patients

Proband II.2, a 13-year-old female patient born to a consanguineous union (**Figure 1A**), first experienced autoinflammatory symptoms when she was 15 months of age. She presented with recurrent episodes of fever as well as oral ulcers occurring once every two weeks. She suffered from abdominal pain and diarrhea during childhood. She presented with asymmetric arthralgia of large joints and persistent folliculitis. Genital ulcers were very rare. Her sister (individual II.1) was 15 years old. Her first symptoms appeared at the age of 5 and they included recurrent episodes of fever, persistent oral ulcers and rare genital ulcers. She also suffered from abdominal pain and diarrhea during childhood. Her skin lesions were acne and folliculitis. Their father (individual I.1) was 38 years old. His first symptoms appeared at the age of 15 and included episodes of fever, oral and genital ulcers occurring once every two weeks. He suffered from abdominal pain with diarrhea twice per year and colonic ulcerations were detected at colonoscopy. He also presented with myalgia and arthralgia of the knees or ankles. His skin lesions included folliculitis. He suffered from episcleritis and uveitis at two occasions. Individuals II.1 and II.2 show significant improvement of their symptoms upon colchicine-treatment. Individual I.1 received colchicine treatment and then tapered courses of oral corticosteroids and azathioprine, leading to a satisfactory long-term control of the disease. The clinical features of the three patients are summarized in table 1.

**Figure 1.**
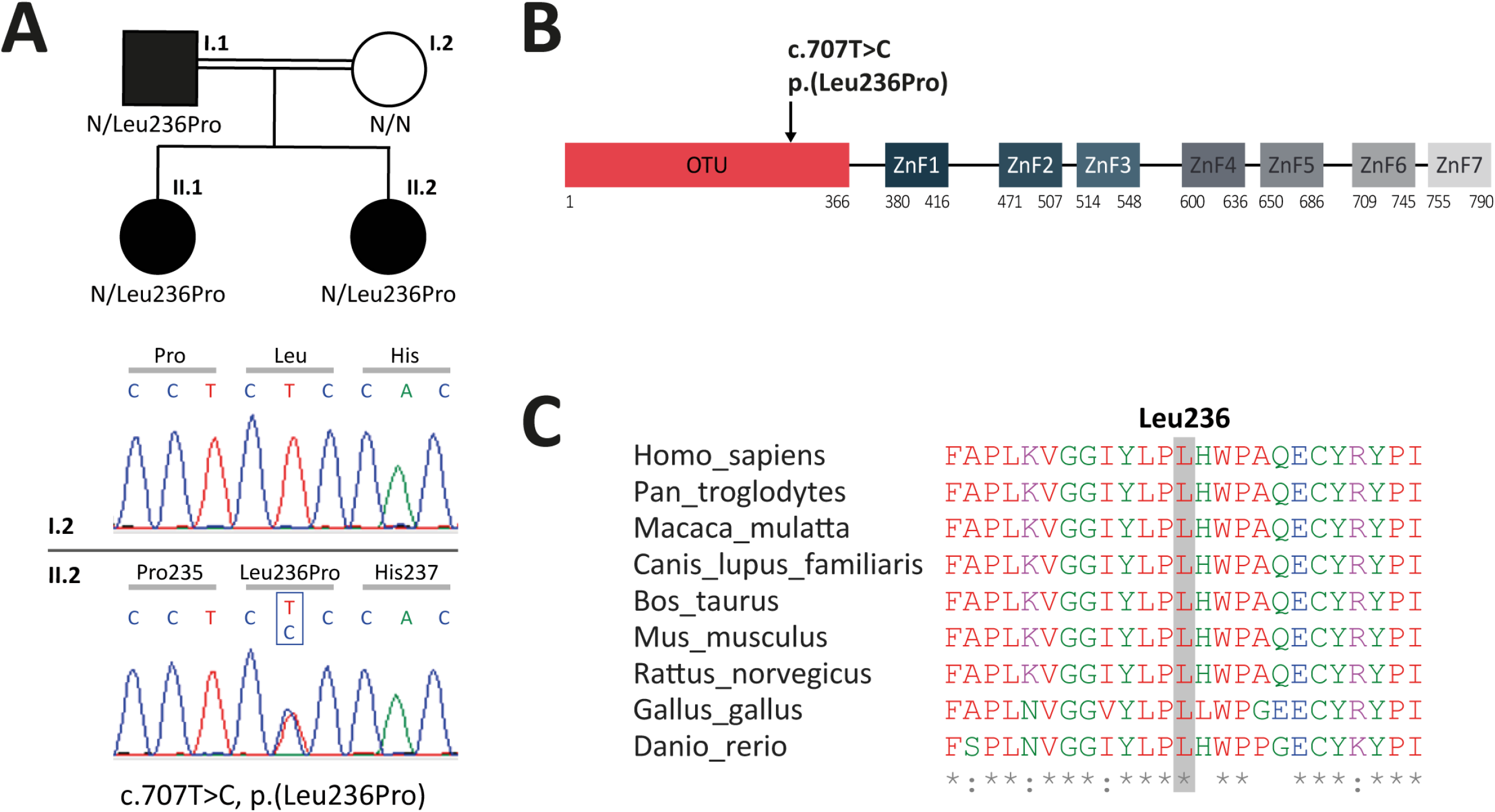
A. Upper panel - Genealogical tree of the patients with HA20 (Individuals I.1, II.1 and II.2). Lower panel - Electropherograms from Sanger sequencing showing the heterozygous c.707T>C, p.(Leu236Pro) variation in individual II.2 and not in her healthy mother (individual I.2). B. Schematic representation of A20 with domain organization showing the OTU domain harboring the p.(Leu236Pro) variation as well as the ZnF domains. C. Evolutionary conservation of the A20 Leu236 residue across species.

### Identification of a novel missense variation in *TNFAIP3*

To identify the molecular basis of the patients’ disease, targeted next generation sequencing (NGS) for autoinflammatory diseases was conducted on a DNA sample from proband II.2. It revealed a non-previously reported heterozygous variation c.707T>C, p.(Leu236Pro) in the *TNFAIP3* gene. Sanger sequencing confirmed this result in the proband and identified this variation in the related affected individuals (I.1 and II.1) (**Figure 1A**). The affected leucine residue at position 236 (Leu236) is located in the OTU domain of A20 **(Figure 1B**) and is invariant across species (**Figure 1C**) supporting a deleterious effect of the substitution of this amino acid. The c.707T>C, p.(Leu236Pro) variant has not been described in sequence-variant databases, such as the genome aggregation database (gnomAD, https://gnomad.broadinstitute.org). The Combined Annotation-Dependent Depletion (CADD, https://cadd.gs.washington.edu/snv) tool provides a score of 29.5 for the c.707T>C, p.(Leu236Pro) variation, predictive of a pathogenic variation.

### Cytokine profile in patients-derived PBMCs

In order to determine the cytokine profile of the patients identified in this study, we quantified inflammatory cytokines in plasma samples and in PBMC supernatants upon LPS cell treatment. Plasma levels of pro-inflammatory cytokines (IL1β, IL18, IL6, TNFα, IFNγ, and IP10) were substantially elevated in the patients (**Figure 2** – upper panel), as compared to healthy individuals. The patients also exhibited high levels of the anti-inflammatory cytokine IL10. In LPS-stimulated PBMCs, pro-inflammatory cytokine levels were higher in the patients compared to healthy individuals (**Figure 2** – lower panel) except for IL6 which exhibited high levels in the patients’ samples prior to stimulation. IFNγ, IP10 and IL10 cytokines were not detected in the PBMC supernatants of healthy individuals or HA20 patients.

**Figure 2.**
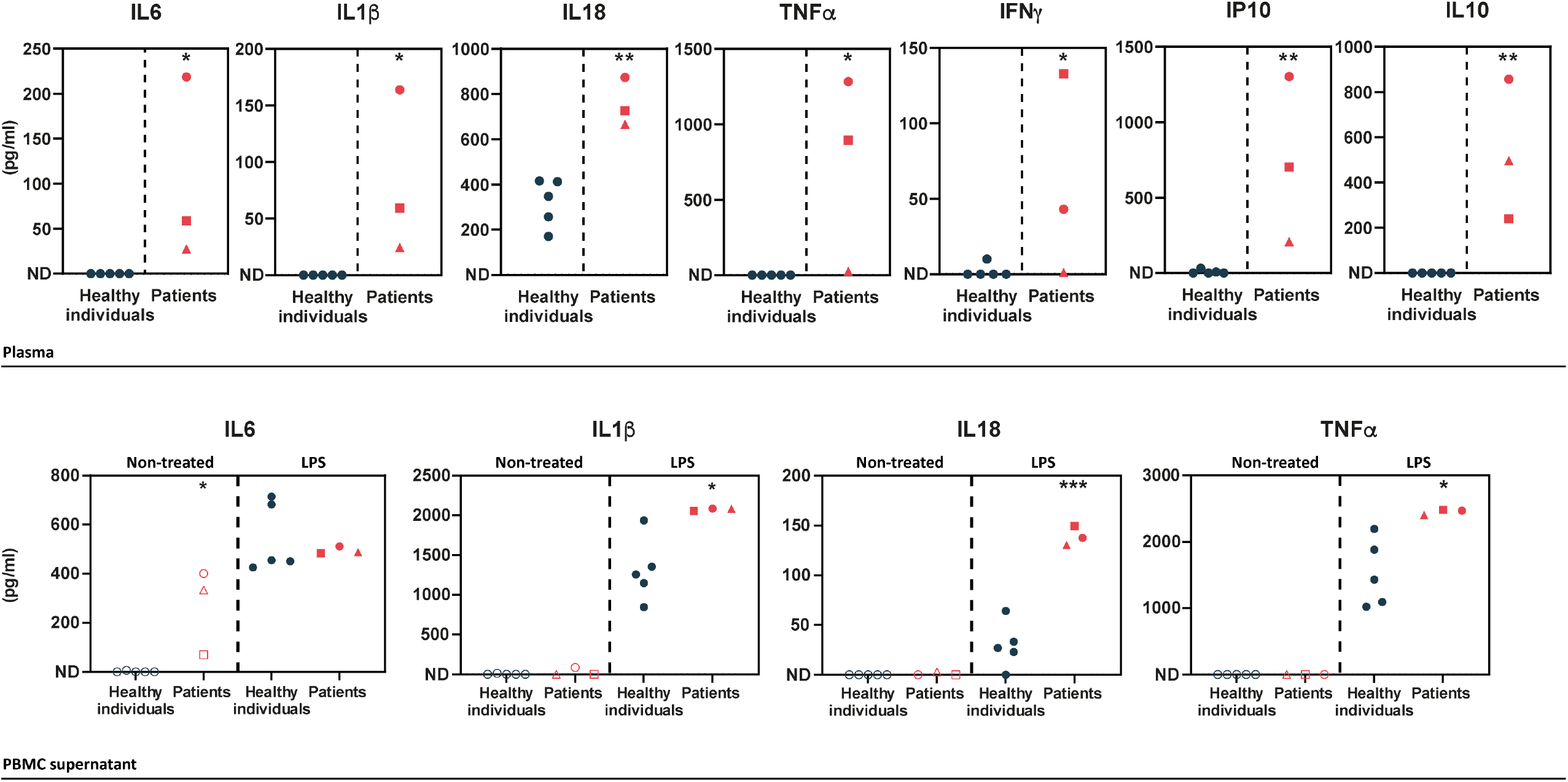
**Upper panel**. ELISA-assessed plasma cytokine levels in the patients (Individual I.1: red square, individual II.1: red triangle and individual II.2: red circle) and 5 healthy donors (blue circles). **Lower panel.** ELISA-assessed cytokine levels in PBMC supernatants prior to (empty symbols) and upon (filled symbols) LPS treatment (100 ng/mL for 16h) (Individual I.1: red square, individual II.1: red triangle and individual II.2 red circle, and 5 healthy donors: blue circles). ND: Not Detected. A two-tailed t-test was used and asterisks indicate that the mean of the cytokine levels quantified in samples from five healthy individuals is significantly different from the mean of the cytokine levels quantified in samples from the three patients. P values <0.05 were considered statistically significant. P values <0.05, <0.01, and <0.001 are indicated with *, **, and *** respectively.

### Evidence for A20 haploinsufficiency from patient-derived samples

To assess the pathogenicity of the p.(Leu236Pro) variation, we first sought to analyze A20 protein expression in patients’ PBMCs. In contrast to two healthy individuals, all three patients (individuals I.1, II.1 and II.2) exhibited a significant reduction of A20 expression levels (**Figure 3A**).

**Figure 3.**
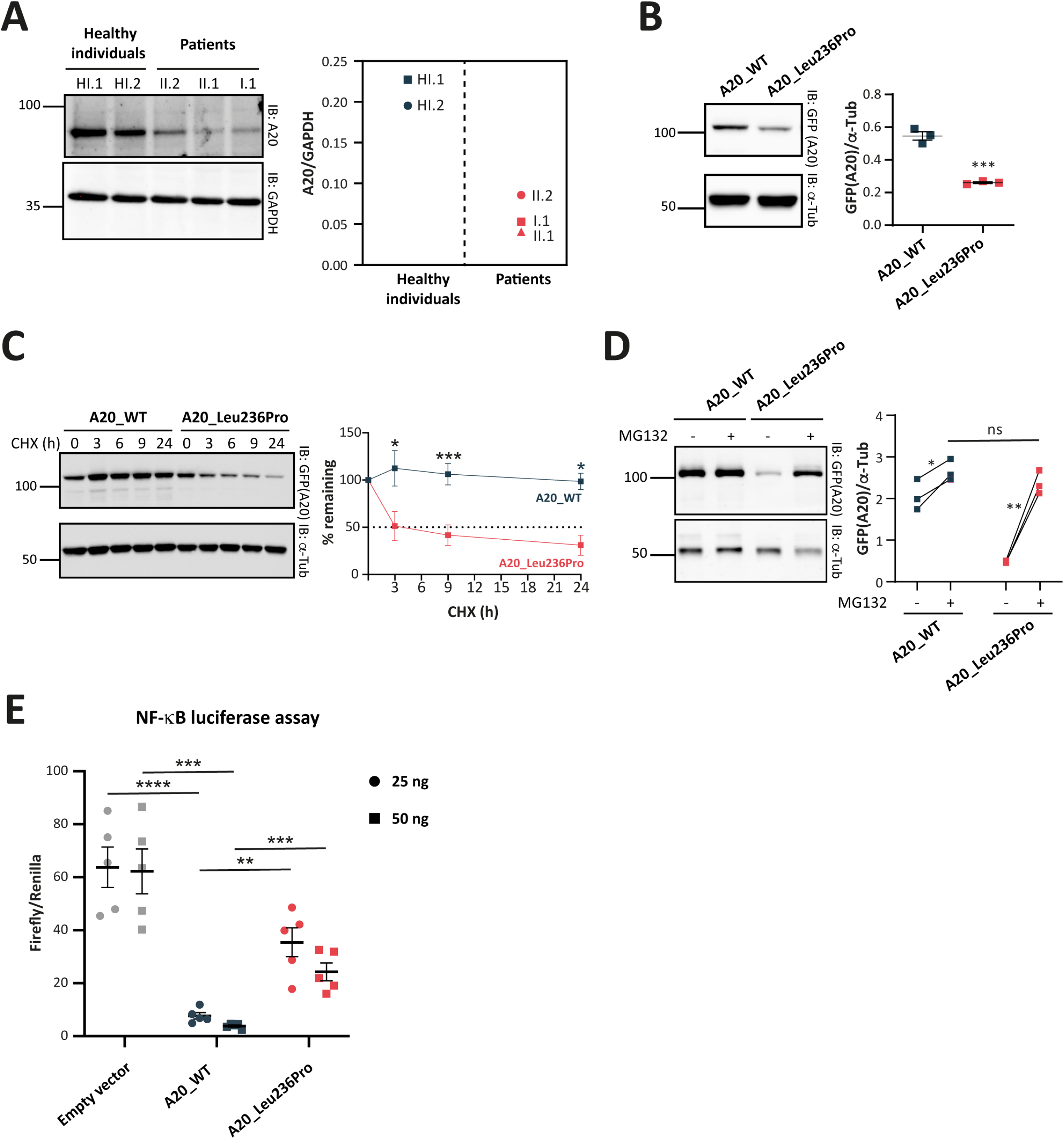
**A.** A20 protein expression in PBMCs of healthy individuals and individuals I.1, II.1 and II.2. **B.** Steady-state protein levels of A20_WT and A20_Leu236Pro upon transient expression in HEK293T (n=3, P value = 0.0004). **C.** Protein levels of A20_WTandA20_Leu236Pro upon transient expression in HEK293T and treatment with cycloheximide (100 μg/ml) at the indicated time points. (n=3, P value = 0.0158, 0.0008 and 0.026 for time points 3, 9 and 24 hours, respectively). **D.** Protein levels of A20_WT and A20_Leu236Pro upon transient expression in HEK293T and treatment with MG132 (20 μM – 5 hours) (n=3, P values for (A20_WT-DMSO vs -MG132), (A20_Leu236Pro-DMSO vs -MG132) and (A20_WT-MG132 vs A20_Leu236Pro-MG132) = 0.037, 0.006 and 0.125, respectively). For all western blot experiments, equal amounts of protein extracts were subjected to SDS-PAGE and immunoblotted with the indicated antibodies. The A20 or GFP (the TNFAIP3-expressing plasmid used is pEGFP-C1-A20) signal was quantified with ImageJ software and normalized to the amount of GAPDH or α-tubulin used as a loading control. **E.** Quantification of the NF-κB signaling in HEK293T cells transiently expressing 25 or 50 ng of empty vector, A20_WT or A20_Leu236Pro. Cells were treated with 10 ng/mL TNFa for 5 hours. Firefly luciferase activity was normalized to the Renilla signal (n=5, P values for (A20_ WT vs Empty vector – 25 ng), (A20_ WT vs A20_Leu236Pro – 25 ng), (A20_ WT vs Empty vector – 50 ng), (A20_WT vs A20_Leu236Pro – 50 ng) are <0.0001, = 0.0011, = 0.0001 and = 0.0003, respectively). For all experiments, two-tailed t-test was used and asterisks indicate that the mean is significantly different; P values <0.05 were considered statistically significant. P values <0.05, <0.01, <0.001, and <0.0001 are indicated with *, **, ***, and ****, respectively.

### Decreased stability of the A20_Leu236Pro protein

With the aim of determining the molecular basis of reduced A20 protein expression in the patients and further assessing the impact of the p.(Leu236Pro) missense variation on the protein, we performed *in silico* stability analyses of the available human crystal structure of the OTU domain of A20 in its active form (Protein Data Bank: 3ZJD) (Kulathu et al., 2013) using the bioinformatics tool PremPS (Yuting Chen et al., 2020). The single variation Leu236Pro of A20 is destabilizing, as indicated by the positive value of 1.820 kcal mol^-1^ for the change of unfolding Gibbs free energy ΔΔG. Similar results were obtained using two other prediction tools: Site Directed Mutator (ΔΔG= 1.860 kcal mol^-1^) and MAESTRO (Multi AgEnt STability pRedictiOn) (ΔΔG= 1.752 kcal mol^-1^).

To confirm these *in silico* results, we assessed the expression and stability of the A20_Leu236Pro protein in HEK293T cells transiently expressing A20_WT and A20_Leu236Pro. As shown in **Figure 3B**, the steady-state protein amounts of A20_Leu236Pro were only 47.6% of that of the WT protein (0.54 for A20_WT vs 0.26 for A20_Leu236Pro). To determine if this difference resulted from protein instability, we compared the half-life of each protein upon inhibition of protein synthesis with cycloheximide (CHX) in HEK293T cells. Quantification of protein amounts at different time points showed that the half-life of A20_Leu236Pro was 2.8 hours, whereas the A20_WT protein was still highly expressed even after 24 hours of treatment (**Figure 3C**). This substantial reduction of the protein half-life shows that A20_Leu236Pro is less stable than its WT counterpart. To test whether this reduced stability was associated with abnormal protein degradation by the ubiquitin-proteasome system (UPS), we inhibited the proteasome-dependent degradation with the peptide-aldehyde inhibitor MG132 in HEK293T cells. As shown in **Figure 3D**, substantial accumulation of A20_Leu236Pro was observed upon inhibition of the proteasome up to 91.3% of the WT protein levels (2.52 for A20_WT vs 2.30 for A20_Leu236Pro). This strongly suggests that the difference of the amounts of WT and Leu236Pro A20 proteins in non-treated cells results from enhanced UPS-dependent degradation.

### Decreased inhibition of NF-κB activity by the A20_Leu236Pro protein

Given the role of A20 in the regulation of the NF-κB activity, we assessed the impact of the p.(Leu236Pro) variation on the ability of A20 to inhibit the canonical NF-κB pathway. To this end, we transiently transfected HEK293T cells with an empty vector, A20_WT or A20_Leu236Pro encoding plasmids along with NF-κB reporter vector and Renilla vector for normalization, and then quantified the luminescence signal upon TNFα treatment (10 ng/mL, 5 hours). Compared to an empty vector, A20_WT exhibited up to 8-fold decrease of the NF-κB activity in a dose-dependent manner (**Figure 3E**). A20_Leu236Pro failed to suppress TNFα-induced NF-κB activity up to the levels of the WT counterpart: at 25 ng, A20_Leu236Pro decreased the NF-κb activity only by 2 folds compared to the empty vector. Moreover, despite increasing the doses of A20_Leu236Pro (50 ng), the level of NF-κB inhibition observed upon A20_WT expression was not reached (**Figure 3E**).

### Identification of an OTU subdomain containing three *TNFAIP3* pathogenic missense variations

To better assess the impact of the localization of *TNFAIP3* missense variations on A20 protein function we reviewed all missense variations reported to date, including the variation herein identified (**Table 2**). The majority (5/8) are located in the OTU domain: c.707T>C, p.(Leu236Pro) (this study), c.305A>G, p.(Asn102Ser) (Yu Chen et al., 2020), c.574G>A, p.(Glu192Lys) (Kadowaki et al., 2021), c.728G>A, p.(Cys243Tyr) (Shigemura et al., 2016) and c.929T>C, p.(Ile310Thr) (Kadowaki et al., 2021). The missense variations c.1428G>A, p.(Met476Ile) (Dong et al., 2019), c.1939A>C, p.(Thr647Pro) (Mulhern et al., 2019) and c.2126A>G, p.(Gln709Arg) (Kadowaki et al., 2021) are located in ZnF2, between Znf4 and ZnF5 and in ZnF6, respectively (**Table 2**).

Among the OTU domain missense variations, the pathogenic significance is demonstrated for three: p.(Leu236Pro) (reported in this study), p.(Glu192Lys) (Kadowaki et al., 2021) and p.(Cys243Tyr) (Shigemura et al., 2016), which all fulfill the American College of Medical Genetics and Genomics (ACMG) criteria (Richards et al., 2015). The remaining two missense variations located in the OTU domain, p.(Asn102Ser) and p.(Ile310Thr), cannot be classified as pathogenic. In fact, the p.(Asn102Ser) variation has an allele frequency in the general population gnomAD database of 3482/281824, which is incompatible with the rare occurrence of HA20; we therefore classified this variation as not pathogenic despite the potential localization of Asn102 residue within a pocket in the active site of A20 that is important for its deubiquitinase activity (Komander and Barford, 2008; Lin et al., 2008). As for the p.(Ile310Thr) variation, it was qualified by the authors of uncertain significance due to the lack of evolutionary conservation of the affected amino acid together with the absence of functional alteration of the variant (Kadowaki et al., 2021). Among the three *TNFAIP3* missense variations located outside the OTU domain, only the p.(Met476Ile) variation can be considered as pathogenic. The other two variations, c.1939A>C, p.(Thr647Pro) and c.2126A>G, p.(Gln709Arg), are not in favor of HA20 diagnosis. Information about the pathogenic significance of the c.1939A>C, p.(Thr647Pro) variation is conflicting; despite disruption of the inhibitory effect of this variant on NF-κB reporter gene activity (Kadowaki et al., 2021), its allele frequency in gnomAD is 511/282864 and the individuals carrying this variation lack HA20 hallmark clinical features (Mulhern et al., 2019) (**Table 2**). As for the c.2126A>G, p.(Gln709Arg) variation, it is considered as likely benign by the authors due to the lack of evolutionary conservation of Gln709 and the absence of protein function alteration (Kadowaki et al., 2021).

We further studied the impact of the three pathogenic missense variations located in the OTU domain on the 3D-structure of A20. Notably, the positioning of Leu236, Glu192 and Cys243 residues on the human crystal of A20 OTU domain revealed that Glu192 and Cys243 are in close vicinity to Leu236, suggesting possible common interactions (**Figure 4A**). To address this hypothesis, we first used the PremPS prediction tool to model the interactions established by each of these residues within the WT protein and to predict the impact of each variation on these interactions (**Figures 4B, 4C** and **4D**). In A20_WT, Leu236 establishes several hydrogen and/or polar bonds as well as hydrophobic interactions with the surrounding residues; some of these interactions are altered in A20_Leu236Pro, e.g., interactions with Phe197, Asn201, and Tyr306 (**Figures 4B**). Modified interactions are also observed in A20_Glu192Lys where Lys192 establishes additional interactions with Leu171 and Met174 compared to Glu192 (**Figure 4C**) and particularly in A20_Cys243Tyr where the phenol group of Tyr243 induces substantial change of amino acid interactions (**Figure 4D**). Additionally, we sought to identify overlapping interaction alterations between these three variations (**Figure 4E**). Indeed, when comparing the alterations of amino acid interactionscaused by both p.(Leu236Pro) and p.(Glu192Lys), we observe that more robust interactions exist between Pro236 and the cycle of Phe197 as compared to the Leu236 in the WT counterpart (**Figure 4B** and **4E**); these altered interactions could impact those established between Phe197 and surrounding residues, notably in the region 194 to 196 that are close to and interact with Glu192 (**Figures 4C, 4E** and **4F**). As for the comparison of the disruptions induced by p.(Leu236Pro) and p.(Cys243Tyr), we first observe that the interactions with residues Asn201 and Tyr306 are lost in A20_Leu236Pro as compared to A20_WT (**Figures 4B** and **4E**) and that interestingly, both Asn201 and Tyr306 do not interact with the WT Cys243 but with variant Tyr243 (**Figures 4D** and **4E**). Similarly, Trp238 that interacts with Leu236 or Pro236 can establish interactions with Tyr243 but not the WT Cys243 (**Figures 4B, 4D** and **4E**). Lastly, since the amino acid in position 235 is a proline, the p.(Leu236Pro) variation results in two consecutive proline residues, which is uncommonly encountered in proteins compared to singlet prolines (Morgan and Rubenstein, 2013) and could hinder proper interactions in this region. For instance, the proper interaction established between Cys243 and Pro235 could be altered by the presence of a proline residue in position 236 as shown in **Figures 4D** and **4E**.

**Figure 4.**
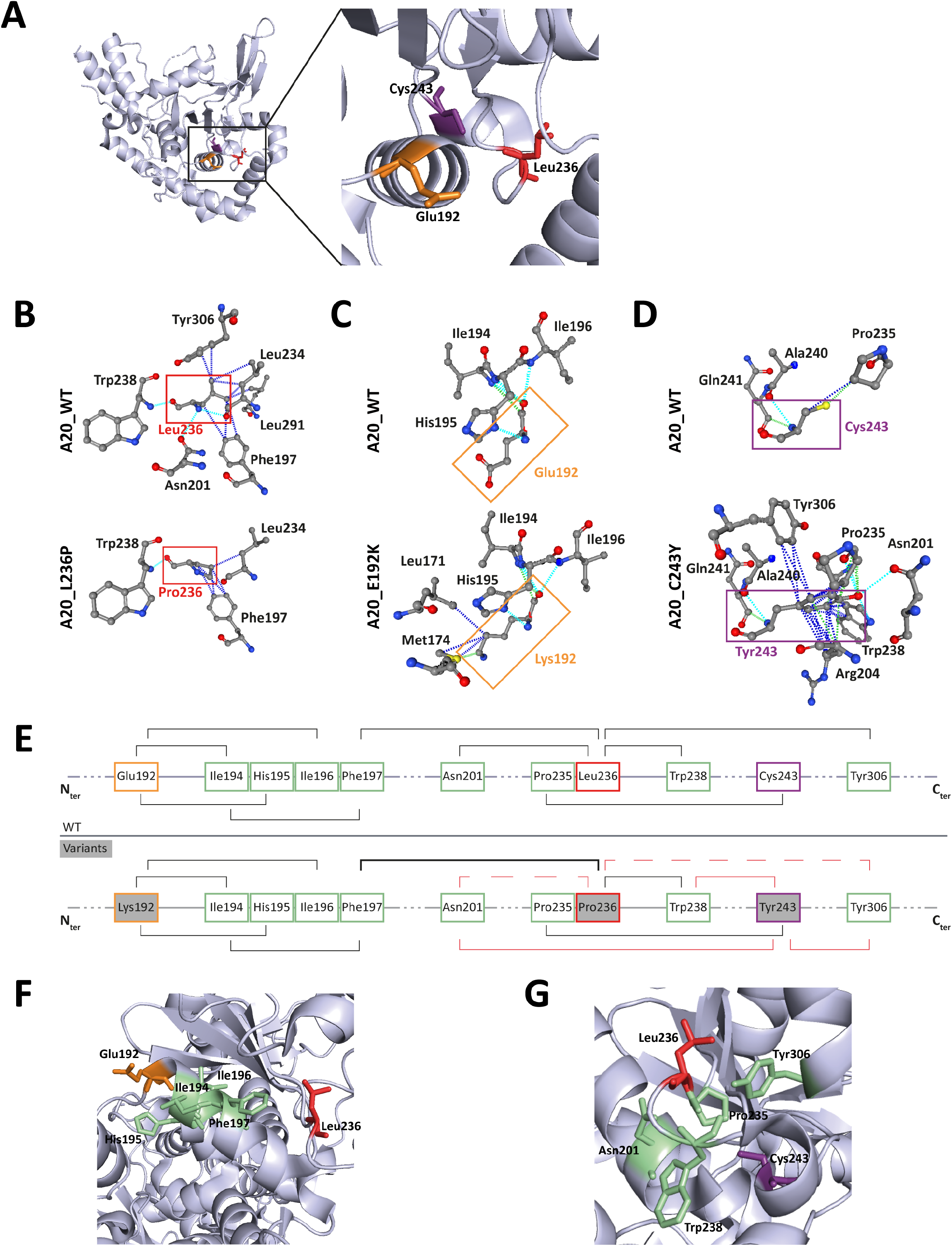
**A.** Amino acids Leu236 (red), Glu192 (orange) and Cys243 (purple) are placed on the crystal structure of the OTU domain of A20 using the 3zjd crystal available of PDB and PyMol software. **B. C. D.** Prediction of the amino acid interactions in A20_WT or A20 variant proteins) established by B: Leu236 (upper panel) or Pro236 (lower panel), C: Glu192 (upper panel) or Lys192 (lower panel), D. C243Y (upper panel) or Tyr243 (lower panel)) using the 3zjd crystal available of PDB and the PremPS in silico prediction tool. Oxygen atoms are represented in red; nitrogen atoms are represented in blue; the remaining atoms are in grey. The interactions are shown in dotted lines: hydrophobic (dark blue), polar (turquoise blue) and Van der Waals (green). **E.** Schematic representation of the predicted amino acid interactions in A20_WT (upper panel) and the variant protein (lower protein). The amino acids involved in pathogenic variations are: Glu192 (orange frame), Leu236 (red frame) and Cys243 (purple frame). On the lower panel, the variant counterpart is represented in grey background and changes in amino acid interactions compared to A20_WT are represented in red: full-lines represent newly formed interaction, dotted lines represent the loss of an interaction compared to A20_WT, the thicker line represents a higher number of possible interactions**. F. G.** Amino acids are placed on the crystal structure of the OTU domain of A20 using the 3zjd crystal available of PDB and PyMol software: residues not affected by variations are shown in light green, Glu192 is shown in orange, Leu236 in red and Cys243 in purple. In F, the helix formed by amino acids 111-149 is hidden to allow for better graphical representation of the amino acids of interest.

This model suggests the existence, on the properly folded protein, of a critical region of the OTU domain encompassing amino acids affected by pathogenic variations and that are distant on the primary sequence: Glu192, Leu236 and Cys243 (**Figure 4G**).

## DISCUSSION

The diagnosis of HA20 is highly challenging due to its heterogeneous clinical presentation and the lack of pathognomonic symptoms. The non-ambiguous pathogenic significance of *TNFAIP3* variations is necessary for the confirmation of HA20 diagnosis and adaptation of the clinical care. While in the case of *TNFAIP3* nonsense or frameshift variations, the deleterious impact is most often established, the pathogenic effect of missense variations is difficult to determine.

Herein, through the in-depth study of three patients from a family carrying a novel *TNFAIP3* missense variation, we determine the pathogenic significance of the c.707T>C, p.(Leu236Pro) variation based on well-established criteria defined by the ACMG (Richards et al., 2015). We provide solid proof for the pathogenicity of this variation relying on the analysis of the clinical and biological features of the patients, the intrafamilial segregation of the variation with the disease, the absence of occurrence of the variation in the general population, the conservation of the affected residue as well as the functional evaluation of the variant, i.e., expression and stability of the variant protein and its ability to inhibit the NF-κB pathway. By assessing two other missense variations in the OTU domain of A20 that fulfill the same criteria, we identify a critical region in the OTU domain in which residues involved in pathogenic missense variations establish common interactions.

The clinical features and the cytokine profile of the patients presented in our study are in line with those exhibited by previously reported HA20 patients. Indeed, elevated levels of pro-inflammatory cytokines in the plasma and supernatants of LPS-stimulated PBMCs of the patients presented in our study resemble that of previously reported patients harboring *TNFAIP3* pathogenic truncating variations (Rajamäki et al., 2018; Zhou et al., 2016). In keeping with this, the patients reported herein exhibit HA20 characteristic features: early-onset, recurrent fever, articular symptoms, recurrent oral, genital and/or gastrointestinal manifestations. Nevertheless, HA20 clinical manifestations can vary considerably between patients even among those who carry the same variation (Aeschlimann et al., 2018). Depending on disease severity, colchicine treatment, corticosteroids or cytokine antagonists can be required for HA20 treatment. Colchicine treatment was satisfactory for the control of the symptoms of individuals II.1 and II.2, suggesting a less severe disease presentation, whereas corticosteroids and azathioprine were required for individual I.1. The difference in disease severity observed in these patients cannot be attributed to the missense nature of the variation. In fact, other HA20 patients carrying pathogenic missense variations were shown to be resistant to colchicine and required steroid therapy (Dong et al., 2019; Kadowaki et al., 2021; Shigemura et al., 2016). Disease management of the patients reported in our study did not require cytokine antagonists. Nevertheless, in light of the elevated IL6 levels in PBMCs supernatant prior to LPS-stimulation, the use of IL6 antagonist for the control of the patients’ symptoms could be of therapeutic interest in case the patients develop resistance to their current treatments, as previously reported for the treatment of two HA20 patients (Lawless et al., 2018; Zhou et al., 2016).

Further findings supporting the pathogenic significance of the p.(Leu236Pro) variation and the similarity of its consequences with other *TNFAIP3* loss-of-function variations include enhanced UPS-dependent degradation of A20_Leu236Pro protein, which is most likely responsible for A20 haploinsufficiency observed in the patients’ PBMCs. Noteworthy, the pathogenic Cys243Tyr variant also exhibits decreased protein levels (Kadowaki et al., 2021). As shown in our study, Leu236 and Cys243 establish common interactions within a small region of A20 suggesting that variations in this region would lead to enhanced recognition by the protein quality control machinery, thus increased degradation. Moreover, our experiments indicate that in addition to reduced levels of the variant protein, the function of A20_Leu236Pro is altered. Indeed, in an attempt to bypass the decreased protein expression *in vitro,* we doubled the amounts of mutant A20 plasmid used for transfection in the NF-κB luciferase assay; although this allowed for an increased inhibition of the canonical NF-κB pathway, the inhibitory function of A20_Leu236Pro was not restored up to the levels of the WT protein. Interestingly, this functional deregulation has also been observed for the pathogenic Cys243Tyr and Glu192Lys variants: a 10-fold protein expression increase was required to compensate the loss-of-function of the Cys243Tyr variant (Shigemura et al., 2016), as for the Glu192Lys variant, it does not exhibit reduced expression compared to the WT protein but it loses the ability to sufficiently inhibit the NF-κB pathway (Kadowaki et al., 2021), in line with its location at the putative ubiquitin-binding site of A20 (Lin et al., 2008).

The consequences on the native structure of A20 of the three pathogenic missense variations of the OTU domain (p.(Leu236Pro), p.(Glu192Lys) and p.(Cys243Tyr)) converge. Although Glu192 and Cys243 are distant, Leu236 bridges the interactions between the two regions they belong to on the tertiary structure of A20. Moreover, p.(Leu236Pro) and p.(Cys243Tyr) have common consequences on interactions established by surrounding amino acids such as Asn201, Pro235, Trp238 and Tyr306, suggesting a putative importance for these residues for proper protein expression and or function.

The OTU domain of A20 harbors the active site essential for the deubiquitinase activity of A20, which encompasses the catalytic Cys103, His256 and a third residue for which Asp70 or Thr97 are possible candidates (Komander and Barford, 2008; Lin et al., 2008). Study of the crystal structure of the A20 OTU domain shows that the ubiquitin binding site of A20 may require the surface formed by α-helices (α4, α7 and α8) and β sheets (β2, β3, β4 and β5) and the loops between them; Glu192 is among the surface residues identified in the ubiquitin binding region, whereas Leu236 and Cys243 are in the loop between β3 and β4 (Lin et al., 2008). Ubiquitin binding requires interactions between hydrophobic residues of ubiquitin and hydrophobic A20 amino acid, making Leu236 a very suitable candidate for surface amino acids in the ubiquitin binding domain. Overall, the HA20-causing missense variations in the OTU domain, which alter the proper function and/or orexpression of the protein, are located in a critical region outside the deubiquitinase active site but close to the ubiquitin binding domain.

## CONCLUSION

In the present study, we analyzed a family with three patients presenting with a phenotype reminiscent of HA20 in which we identified the p.(Leu236Pro) missense variation in the *TNFAIP3* gene that segregates with the disease. We provide evidence from patient-derived samples for the A20 haploinsufficiency, as well as cellular evidence for the loss-of-function effect induced by the variation. By studying the pathogenic missense *TNFAIP3* variations located in the OTU domain of the protein we shed the light on a non-previously identified pathogenic missense variation-containing region. Our data can help in the interpretation of *TNFAIP3* missense variations and in providing an adapted clinical care for the patients.

## PATIENTS AND METHODS

### Affected and healthy individuals

Clinical features were collected through a standardized form. Written informed consents were obtained from each individual for genetic tests according to French legislation and the principles of the Declaration of Helsinki regarding ethical principles for medical research.

### Next generation sequencing

Next-generation sequencing (NGS) was performed using a custom sequence capture (Nimblegen SeqCap EZ Choice system; Roche Sequencing, Pleasanton, California) of the exons and the flanking intronic sequences of the main autoinflammatory disease-causing genes. Sequencing was performed on Nextseq500 platform (Illumina, San Diego, California) according to the manufacturer’s instructions. Variations and segregation analysis were confirmed by standard PCR and Sanger sequencing of DNA extracted from peripheral blood.

### Sanger sequencing

*TNFAIP3* was amplified by PCR and sequenced using the Big-Dye Terminator sequencing kit (Applied Biosystems, Foster City, California) on an ABI3730XL automated capillary DNA sequencer (Applied Biosystems). Sequences were analyzed against the reference sequence (NM_ 006290) using the SeqScape software (Applied Biosystems).

### Plasma and PBMC isolation

Whole blood samples from patients and healthy individuals (provided by the Etablissement Français du Sang) were collected in EDTA-tubes. Plasma was isolated by blood centrifugation at 950g for 10 minutes and kept at −80°C for cytokine measurements. For PBMC (Peripheral Blood Mononuclear Cell) isolation, blood samples were diluted with an equal volume of PBS-EDTA (1 mM) and PBMCs were collected by centrifugation (800g, 15 minutes) of the diluted samples on Pancoll gradient tubes (PANbiotech, Aidenbach). PBMCs were washed three times with PBS-EDTA and incubated at 37°C for 1 hour in RPMI medium before being treated with LPS (100 ng/mL for 16 hours) before proceeding to ELISA cytokine quantification.

### ELISA cytokine quantification

IL1β, IL18, IL6, TNFα, IFNγ, IP10 and IL10 quantification was carried out on plasma and PBMC culture medium using R&D systems kits (Minneapolis, Minnesota) according to the manufacturer’s instructions. For each cytokine quantification, samples from at least five healthy individuals were used as controls.

### Plasmids

Full-length human *TNFAIP3* was purchased from Addgene (pEGFP-C1-A20 - RRID: Addgene_22141). Directed mutagenesis using the Q5 High-Fidelity DNA Polymerase (New England Biolabs, Ipswich, Massachusetts) was used.

### Cell culture and transfection

HEK293T (Human embryonic kidney 293T) cells were maintained in DMEM containing GlutaMAX (ThermoFisher Scientific, Waltham, Massachusetts) supplemented with 10% fetal bovine serum and 1% penicillin/streptomycin (ThermoFisher Scientific) at 37°C in a 5% CO_2_ atmosphere. Cells were transiently transfected using Fugene HD Transfection Reagent (Promega E231A - Madison, Wisconsin) according to the manufacturer’s instructions.

### Proteasome inhibition

HEK293T cells were transfected with the indicated plasmids and treated 24 hours later with 20 μM MG132 (Sigma-Aldrich M7449, Saint-Louis, Missouri) or the equivalent volume of DMSO (Sigma-Aldrich) for 5 hours. Proteins were then extracted and detergent-soluble and insoluble fractions of protein lysates were analyzed.

### Cycloheximide study

HEK293T cells were transfected with the indicated plasmids, and treated 24 hours later with 100 μg/ml cycloheximide (Sigma-Aldrich C4859) or the equivalent volume of DMSO for the indicated times.

### Western blot analysis

Protein extraction and western blot analysis were carried out as previously described (El Khouri et al., 2021). The primary antibodies used were: polyclonal rabbit A20 (Cell Signaling 5630 – RRID: AB_10698880), anti-GFP-HRP (Cell Signaling 2037 - RRID: AB_1281301), anti-GAPDH-HRP (Cell Signaling 51332 - RRID: AB_2799390) and anti-α-Tubulin-HRP (Cell Signaling 9099 – RRID: AB_10695471). The secondary antibodies used were Anti-Rabbit IgG-HRP, Sigma Aldrich A0545 – RRID: AB_257896. Proteins were detected with Amersham ECL Select Western Blotting Detection Reagent (GE healthcare, Chicago, Illinois) and BioRad ChemiDoc Imaging Systems was used for detection. The ImageJ software was used for signal quantification.

### NF-κB luciferase assay

HEK293T cells were transiently transfected with the indicated quantities of *TNFAIP3* WT and mutant plasmids, 100 ng of pGL4.32[luc2P/NF-κB-RE/Hygro] vector and 10 ng of Renilla Luciferase Control Reporter Vector (pGL4-hRluc), purchased from Promega. 24 hours later, cells were treated with 10 ng/mL TNFα for 5 hours then lysed with Passive Lysis Buffer (Promega). Firefly luciferase activity was measured in duplicate according to the manufacturer’s instructions and normalized against the Renilla signal.

### Crystal structure and amino acid interaction analysis

The PremPS (Predicting the Effects of Mutations on Protein Stability) tool, a computational method that predicts protein stability changes induced by missense variations available as a web server (https://lilab.jysw.suda.edu.cn/research/PremPS/) (Yuting Chen et al., 2020) was used to predict protein stability and to analyze amino acid interactions of the Protein Data Bank-available crystal structure of A20 OTU domain in reduced, active state at 1.87 Å resolution (PDB accession number 3ZJD) (Kulathu et al., 2013). The amino acid interactions were shown as provided by the PremPS online tool. Three-dimensional structure was visualized and amino acids were positioned using the PyMol Software.

### Identification of *TNFAIP3* missense variations reported in the literature

Literature review of *TNFAIP3* variations was performed in PubMed using the terms “HA20”, “A20” or “TNFAIP3” and “variation” or “mutation”.

### Statistical analyses

Statistical analyses and graph representation were performed using GraphPad Prism 9.0 software. Unpaired two-tailed Student’s t tests were used for comparisons of the means of two groups. The means of the data provided by sets of independent experiments carried out on samples from the patients (or cells expressing the variant protein) were compared to the means of data provided by sets of independent experiments carried out on the samples from healthy individuals (or cells expressing the WT protein). The number of replicates (n) corresponds to biological replicates. Data are presented as means ± SEM and individual data points are represented when required. P values <0.05 were considered statistically significant. P values <0.05, <0.01, <0.001, and <0.0001 are indicated with *, **, ***, and ****, respectively.

## DATA AVAILABILITY

Uncropped western blots are provided as Source Data files for Figures 3A, 3B, 3C and 3D.

## ACKNOWLEDGMENTS

We thank the patients and their family as well as the control individuals for their cooperation. The research reported in this manuscript was funded by: Institut National de la Santé et de la Recherche Médicale (INSERM); Agence Nationale de la Recherche (ANR-17-CE17-0021-01); European Union’s Horizon 2020 research and innovation programme under grant agreement No 779295; and Sorbonne Université EMERGENCE-PhenomAID.

The authors declare that they have no conflict of interest.

---

**Figure 3**—*source data 1*

*A20 protein expression in PBMCs of healthy individuals and individuals I.1, II.1 and II.2. Uncropped Western blot images of A20 and GAPDH protein expression in PBMCs of the patients and healthy individuals.*

**Figure 3**—*source data 2*

*Protein levels of A20_WT and A20_Leu236Pro upon transient expression in HEK293T.*

*Uncropped Western blot images of GFP-A20 and α-tubulin protein expression (n=4).*

**Figure 3**—*source data 3*

*Protein levels of A20_WT and A20_Leu236Pro upon transient expression in HEK293T and treatment with cycloheximide.*

*Uncropped Western blot images of GFP-A20 and α-tubulin protein expression (n=3).*

**Figure 3**—*source data 4*

*Protein levels of A20_WTand A20_Leu236Pro upon transient expression in HEK293T and treatment with MG132.*

*Uncropped Western blot images of GFP-A20 and α-tubulin protein expression (n=3).*

## Notes

### Competing Interest Statement

The authors have declared no competing interest.

